# Focal adhesion kinase confers pro-migratory and anti-apoptotic properties and is a potential therapeutic target in Ewing sarcoma

**DOI:** 10.1101/604207

**Authors:** Konrad Steinestel, Esther-Pia Jansen, Marcel Trautmann, Uta Dirksen, Jan Rehkämper, Jan-Henrik Mikesch, Julia S. Gerke, Martin F. Orth, Giuseppina Sannino, Eva Wardelmann, Thomas G. P. Grünewald, Wolfgang Hartmann

## Abstract

Oncogenesis of Ewing sarcoma (EwS), the second most common malignant bone tumor of childhood and adolescence, is dependent on the expression of chimeric EWSR1-ETS fusion oncogenes, most often EWSR1-FLI1 (E/F).

E/F expression leads to dysregulation of focal adhesions (FAs) enhancing the migratory capacity of EwS cells. Here we show that, in EwS cell lines and tissue samples, focal adhesion kinase (FAK) is expressed and phosphorylated at Y397 in an E/F-dependent way involving Ezrin. Employing different EwS cell as *in vitro* models, we found that key malignant properties of E/F are mediated via substrate-independent autophosphorylation of FAK on Y397. This phosphorylation results in enhanced FA formation, Rho-dependent cell migration, and impaired caspase-3-mediated apoptosis *in vitro*. Conversely, treatment with the FAK inhibitor Y15 enhanced caspase-mediated apoptosis and EwS cell migration, independent from the respective EWSR1-ETS fusion type, mimicking an anoikis-like phenotype. Our findings were confirmed *in vivo* using an avian chorioallantoic membrane (CAM) model. Our results provide a first rationale for the therapeutic use of FAK inhibitors to impair metastatic dissemination of EwS.

## INTRODUCTION

Ewing sarcoma (EwS) is a highly aggressive cancer of bone and soft tissue that predominantly affects children and adolescents. Frequently, (micro-)metastasis is already present at the time of diagnosis, and no effective therapy strategies have been established for these patients yet ^6, 33^. Knowledge of the biological mechanisms underlying EwS cell migration might provide rationales for developing urgently needed targeted therapies to prevent or to slow down metastatic dissemination of EwS cells.

EwS are genetically stable tumors characterized by a chromosomal translocation leading to fusion of the *EWSR1* gene on chromosome 22 and variable members of the ETS family of transcription factors. In 85% of cases, the translocation partner for *EWSR1* is *FLI1* on chromosome 11; other possible fusion partners are, among others, the *ERG, FEV* or *ETV1/4* genes ^19, 26^. The resulting chimeric transcription factor *EWSR1-FLI1* (E/F) modulates the expression of a large number of target genes, leading to oncogenic transformation of the cell harboring this fusion ^1^.

So far, only a few studies have investigated the effect of E/F expression on cell migration and invasion of EwS cells as a prerequisite to metastasis. Using an RNA interference approach, Chaturvedi *et al*. demonstrated that knockdown of E/F enhanced tumor cell adhesion and spreading; in a subsequent study, the same group showed that E/F-induced repression of Zyxin and α5 integrin impairs cell adhesion and actin cytoskeletal integrity, thus supporting anchorage-independent cell growth ^3, 4^. CXCR4-dependent migration and invasion of EwS cells were shown to depend on the activity of small Rho-GTPases RAC and CDC42 ^16^.

Focal adhesion kinase (FAK) is a substrate of SRC kinase and localizes to focal adhesions (FAs), where it is activated by integrins, resulting in FA reorganization and increased cell motility ^20^. Integrin-mediated autophosphorylation of FAK on Y397 inhibits detachment-dependent apoptosis, a process named anoikis ^7^. There is experimental evidence that autophosphorylation of FAK on Y397 results from dimerization of two FAK molecules ^32^. Alternatively, FAK autophosphorylation can be induced by the FAK interaction partner Ezrin independent from cell-matrix adhesion ^22^. Ectopic expression of constitutively active FAK rescues cancer cells from induced anoikis ^1^. Moreover, FAK activation appears to be essential for migration as well as mechano-induced osteogenic differentiation of mesenchymal stem cells (MSCs) that are believed to constitute the cells of origin in EwS ^25, 27, 34^. One report previously identified FAK as a potential therapeutic target in EwS by the use of high-throughput tyrosine kinase activity profiling ^5^. In line, another report showed that miR-138, via targeting *FAK*, inhibits proliferation and mobility and induces anoikis of EwS cells ^31^. However, the exact role of FAK in EwS remains elusive.

Here, we show that E/F-dependent autophosphorylation of FAK is a crucial mechanism underlying EwS aggressivity as it promotes a migratory phenotype and inhibits caspase-mediated apoptosis. We show that this mechanism can effectively be targeted by the FAK inhibitor Y15, independent from the respective *EWSR1-ETS* fusion type. Hence, our results point toward a possible use of FAK inhibitors to prevent metastatic dissemination in EwS.

## RESULTS AND DISCUSSION

### Doxycycline (DOX)-inducible down regulation of *EWSR1-FLI-1* (E/F) attenuates Ezrin expression and Y397-phosphorylation of FAK in A673 EwS cells

In order to investigate E/F-dependent changes in FA gene expression levels, a DOX-inducible E/F knockdown in A673 EwS cells was performed, revealing decreased levels of *Ezrin* mRNA upon E/F knockdown (Fig. 1 A). Ezrin has previously been shown to induce FAK autophosphorylation independent from SRC signaling and FAK/integrin interaction^22^. Correspondingly, immunohistochemical analyses of xenograft tumors derived from A673 cells showed significantly decreased Ezrin protein expression and Y397-phosphorylation of FAK upon loss of E/F while total FAK protein expression was unaffected (Fig. 1 B). These findings are in line with previous results showing i) strong expression of Ezrin in primary EwS samples and ii) experimental data supporting a central role of Ezrin in both autophosphorylation of FAK and growth and metastatic spread of EwS^15, 22^.

**Figure 1.**
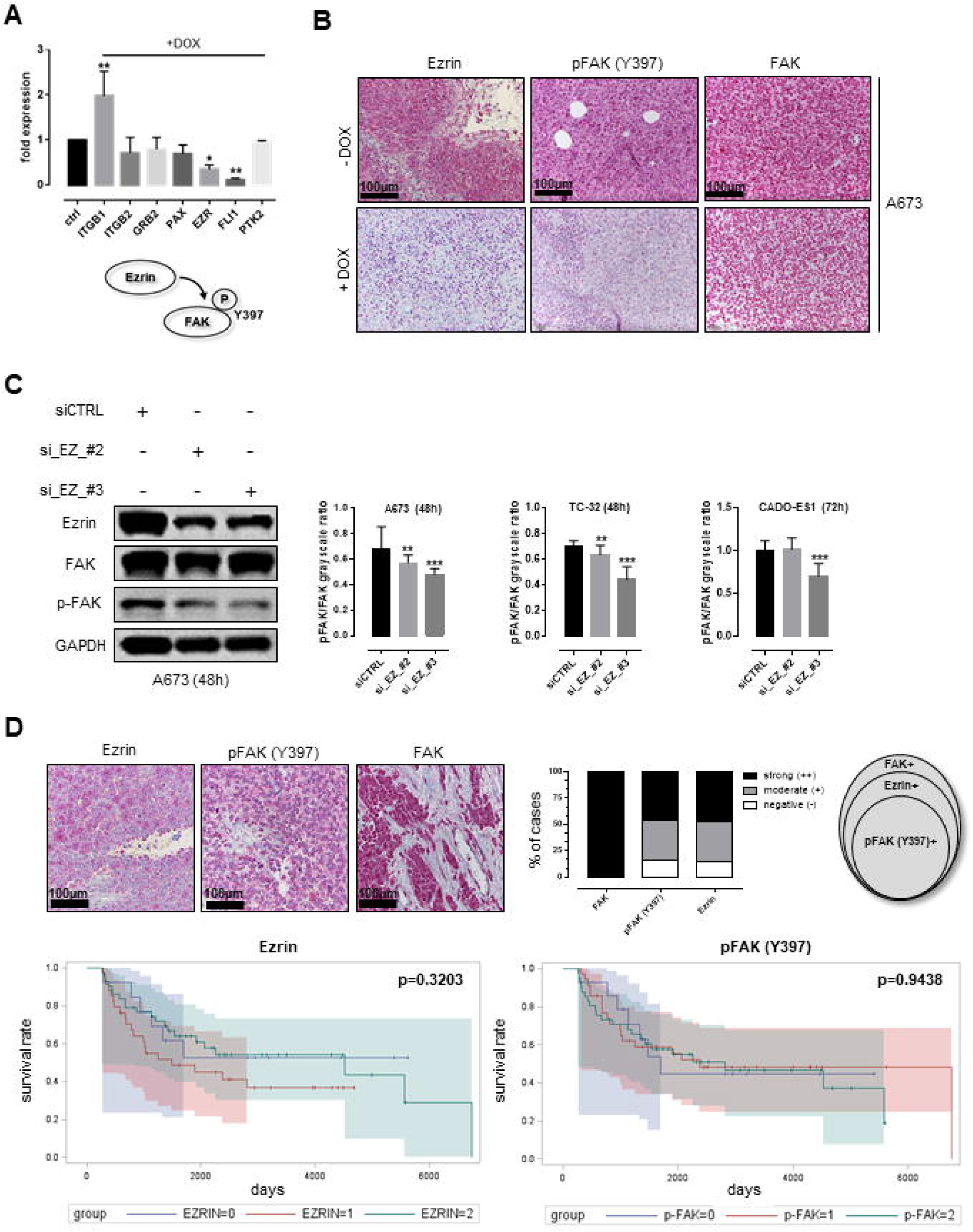
EWSR1-FLI1-dependent Ezrin expression and FAK Y397-phosphorylation in an Ewing sarcoma (EwS) *in vivo* model and primary human tumor samples. **A,** DOX-inducible knockdown of EWSR1-FLI1 in A673 cells showed downregulation of FLI-1 and Ezrin mRNA levels in A673 EwS cells, while FAK levels remained unaffected. **B,** Immunohistochemistry confirmed the downregulation of Ezrin on protein level along with a decrease in Y397-phosphorylation of FAK in A673-derived xenografts while FAK protein levels remained unaltered. **C,** siRNA-based knockdown of Ezrin led to a significant decrease in Y397-phosphorylation of FAK in EwS cells. **D,** Immunohistochemical analyses of FAK/Ezrin expression and Y397-phosphorylation of FAK in tissue samples derived from 97 EwS patients treated in the cooperative Ewing sarcoma study (CESS) trial. All EwS tumor samples were positive for FAK protein in immunohistochemistry, and the majority of tumor samples showed moderate to strong expression of Ezrin and Y397-phosphorylation of FAK (79% and 78% of cases, respectively). All pFAK+ cases were Ezrin+. Survival analyses showed no significant association between Ezrin expression/ FAK phosphorylation and overall survival (*p*=0.3203 and 0.9438, respectively).

### Y397-phosphorylation of FAK in EwS cells depends on the presence of Ezrin

For further *in vitro* studies, we employed the EwS cell lines A673, TC-32 (both EWSR1-FLI1 fusion type 1) and CADO-ES1 (EWSR1-ERG fusion gene) (Supplementary Fig. 1) ^18, 29^. All EwS cell lines strongly expressed Ezrin (Supplementary Fig. 1 E). siRNA-based knockdown of Ezrin significantly impaired Y397-phosphorylation of FAK in A673 (Fig. 1C), TC-32 and CADO-ES1 cells (Supplementary Figure 1 B). Notably, in CADO-ES1 cells, the effect on Y397-phosphorylation was limited to one Ezrin siRNA (#3) and could only be observed 72h after transfection. Taken together, these findings suggest a model in which FAK autophosphorylation in EwS occurs dependent on E/F-driven Ezrin expression.

### FAK is ubiquitously expressed and phosphorylated at Y397 in Ezrin-positive EwS tumor samples

To investigate the relevance of our *in vitro* and *ex vivo* findings in patients, immunohistochemical analyses were performed in a set of tumor samples homogeneously treated in the EWING99 trial (*n*=93; Fig. 1 D; Supplementary Table 1). All investigated samples showed moderate to strong FAK staining with Ezrin being expressed in 79% of tumors. Phosphorylated FAK (Y397) was detected in 78% of samples, which were all simultaneously Ezrin-positive (*p*<0.0001, Chi-square test). Surprisingly, there was no significant correlation between FAK expression levels, Ezrin expression, or phosphorylation of FAK and clinicopathological characteristics, such as response to chemotherapy, first relapse or presence of metastatic disease at the time of diagnosis. This lack of clinical association may well be due to the fact that a vast majority of samples in our cohort were Ezrin/pFAK double-positive (79% and 78%, respectively). However, in concordance with our results, Krishnan *et al*., investigating a similar percentage of strongly Ezrin-positive EwS samples, found no significant differences in the “ezrin score” of patients with localized versus metastatic disease^15^. Thus, the biological impact of Ezrin on the course and outcome of EwS remains to be elucidated as it might be masked by possible alternative signaling pathways in Ezrin-negative EwS.

### EwS cell lines show adhesion- and fusion type-independent Y397-phosphorylation of FAK that can be targeted by 1,2,4,5-Benzenetetraamine tetrahydrochloride (FAK inhibitor 14, Y15)

A673, TC-32 and CADO-ES1 remained viable and showed enhanced FAK phosphorylation when grown under non-adherent conditions (Fig. 2 A), which is of special interest since it had previously been shown that Y397-phosphorylation of FAK inhibits anoikis (detachment-dependent apoptosis)^7^. The compounds PF-573228 (PF-228) and Y15 are potent and specific inhibitors of FAK autophosphorylation activity^9, 28^. Y15 has previously been shown to decrease cell viability and clonogenicity of various carcinomas and to increase detachment, cause apoptosis and inhibit invasion of glioblastoma cells through the inhibition of FAK ^10, 35^. We could confirm that both Y15 and PF-228 (data not shown) abrogate Y397-phosphorylation of FAK in EwS; we selected Y15 for further experiments since its *in vivo* activity has already been documented ^12^. Application of 10 μmol/l Y15 effectively impaired FAK phosphorylation in all three cell lines (Fig. 2 B). The decrease in FAK phosphorylation upon Y15 treatment occurred in parallel with an increase in caspase-3 cleavage and apoptosis as shown by immunoblotting and ApoTox Glo Triplex assays. Fluorescence microscopy and image analysis revealed a significant decrease in the number and size of focal adhesions together with a loss of dorsal actin stress fibers (Fig. 2 C). The decrease in FA formation together with increased caspase 3-mediated apoptosis points towards enhanced anoikis in Y15-treated EwS cells, given that caspase 3 is one of the key effector molecules of detachment-dependent cell death^8^. Moreover, application of 10 μmol/l Y15 significantly impaired cell migration of A673, TC-32 and CADO-ES1 in real-time cell migration assays (Fig. 2 D) together with a decrease in active Rho levels as shown by Rho activity pulldown assays (Fig. 2 E). Since FAK has previously been identified as a potential therapeutic target in EwS, and since migration and invasion of EwS depend on the activity of small Rho-GTPases^5, 16^, these findings support a potential therapeutic role of FAK inhibition in prevention or slowing down metastatic dissemination of EwS.

**Figure 2.**
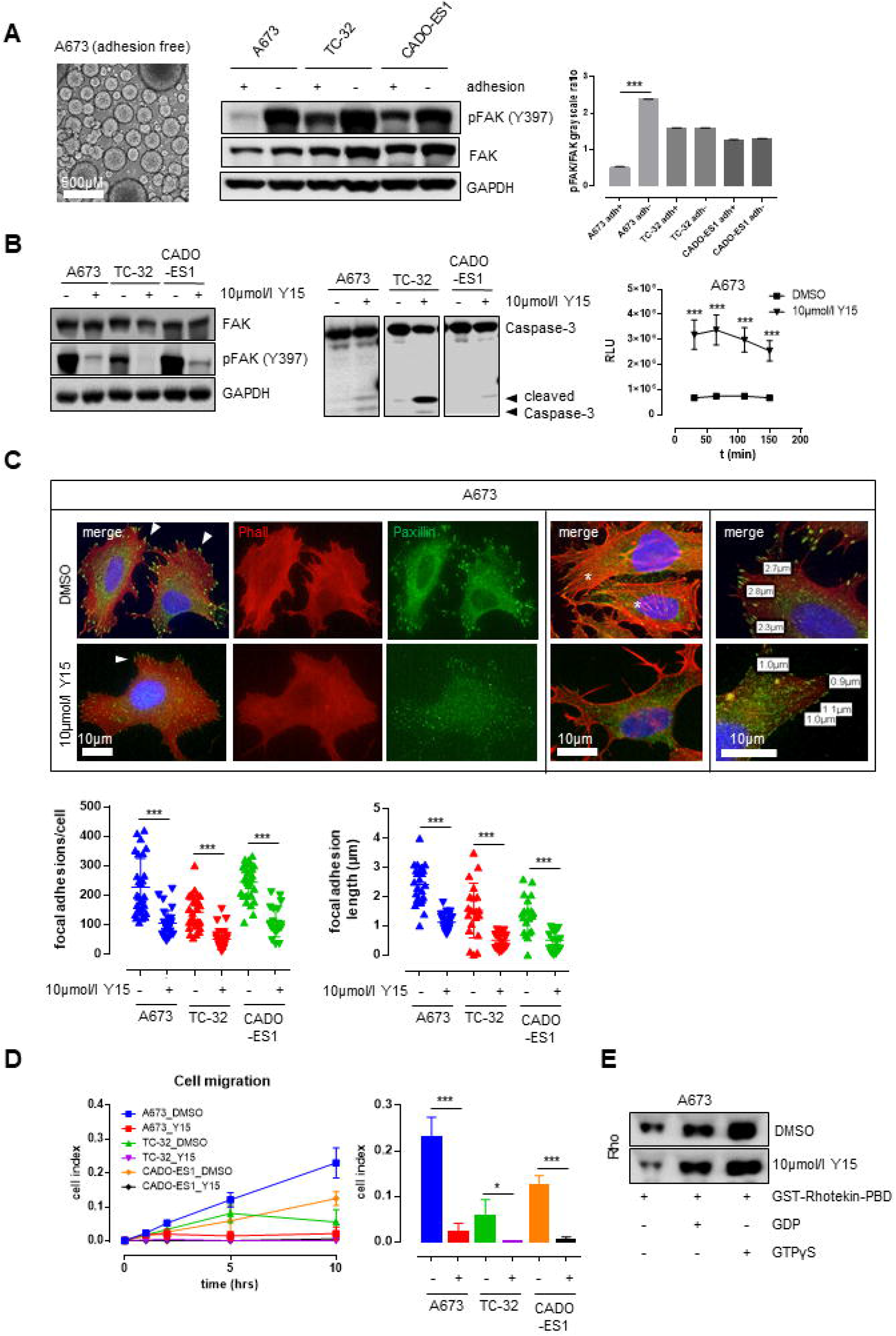
Y397-phosphorylation of FAK under different cell culture conditions and upon application of the FAK inhibitor 1,2,4,5-Benzenetetraamine tetrahydrochloride (Y15) **A,** In EwS cell lines grown under adherent and non-adherent conditions, Y397-phosphorylation of FAK was not only persistent, but increased under adhesion-free conditions in three different EwS cell lines (A673, TC-32 and CADO-ES1) as shown by Western immunoblotting and densitometric analysis. **B,** Application of 10 μmol/l Y15 significantly impaired FAK Y397-phosphorylation and induced cleavage of caspase-3 in in all three EwS cell lines in comparison to DMSO treatment. This was accompanied by a significant increase in apoptosis (ApoTox Glo Triplex assay, shown here are the results from A673 cells). **C,** ImageJ software-based analysis of fluorescence microscopy images showed that treatment of EwS cells with 10 μmol/l Y15 significantly decreased both number (left graph) and size (right graph) of Paxillin-positive focal adhesions (arrowheads) together with a loss of dorsal stress fibers (asterisks) in all three investigated EwS cell lines. **D,** Cell migration was significantly impaired upon treatment with 10 μmol/l of Y15 compared to DMSO in A673, TC-32 and CADO-ES1 cells. **E,** Reduction in migratory capacity of A673 cells upon application of 10 μmol/l Y15 occurred in parallel with a significant decrease in active Rho levels as shown by Rho activity pulldown assay.

### Y15 impairs viability and invasion of EwS in an *in vivo* (avian CAM) model

A chicken chorioallantoic membrane (CAM) model was used to assess the effects of Y15 on EwS tumor xenografts *in vivo*. Application of 10 μmol/l Y15 alone did not have morphologic effects, particularly with regard to the microvascular architecture of the CAM (Fig. 3 A). However, treatment of A673 cells with 10 μmol/l Y15 prior to cell seeding significantly impaired tumor formation (*p*=0.0122) and led to a significant decrease in the size of invasive experimental EwS (*p*=0.0095) along with widespread tumor regression, necrosis and microcalcification with only few residual tumor cell clusters (Fig. 3 B). Immunohistochemistry showed expression of FAK in both, Y15-treated and control cells, but barely detectable Y397-phosphorylation of FAK in the few residual tumor cells after Y15-treatment (*p*<0.001).

**Figure 3.**
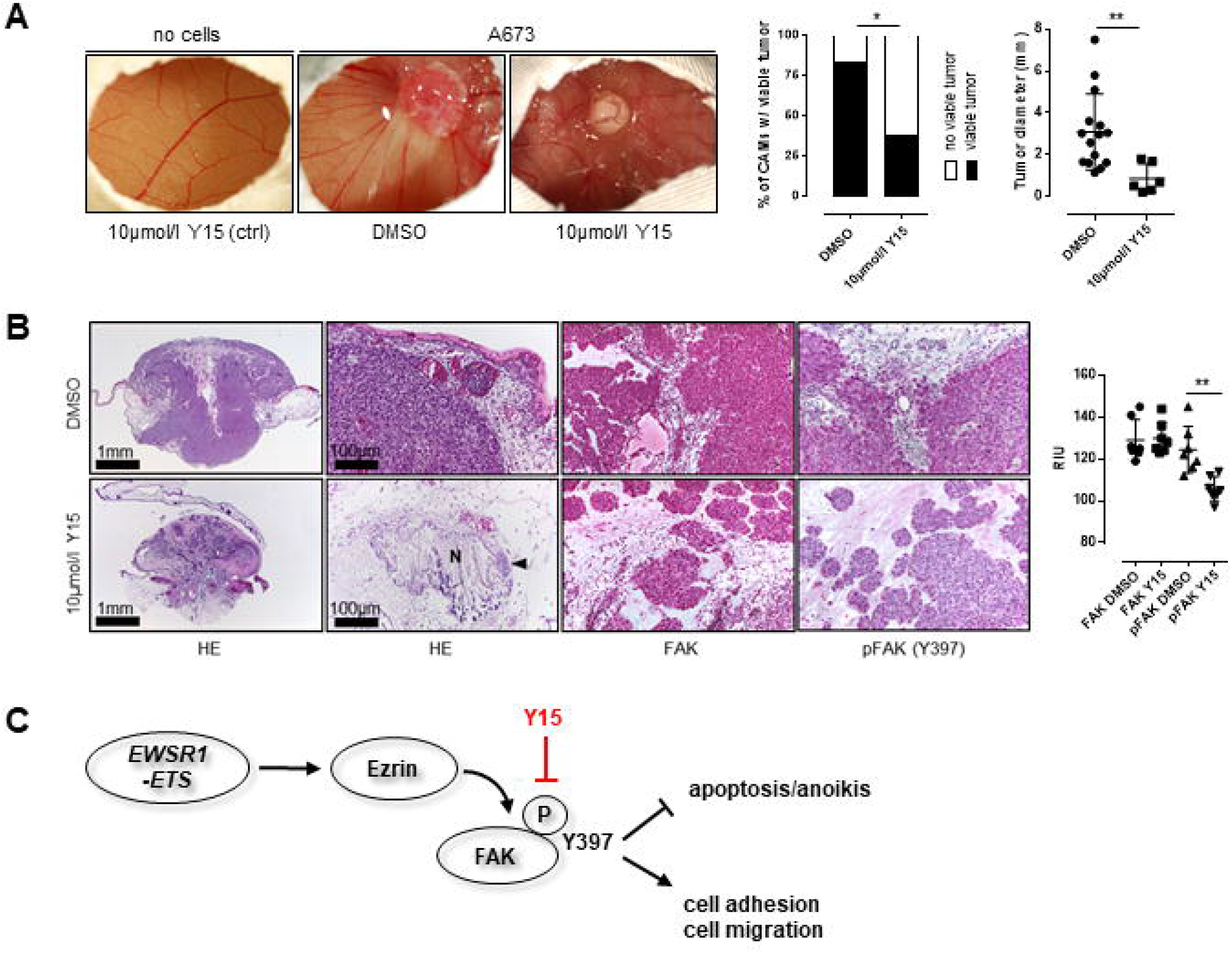
Effects of Y15 treatment on EwS viability and invasion in an *in vivo* (avian CAM) model. **A,** Application of 10 μmol/l Y15 to A673 EwS cells 24h prior to cell seeding significantly impaired the rate of tumor formation as well as tumor size in the CAM model (*p*=0.0122 and *p*=0.0095, respectively) while not altering the (micro-)vascular architecture of the CAM. **B,** Histological analyses showed tumor regression, necrosis (**N**) and microcalcification with only very few residual tumor cells detectable (**arrowhead**) upon Y15 treatment. Immunohistochemistry and quantification via reciprocal intensity showed persistent expression of FAK in both, Y15-treated and control cells, but a significant decrease of FAK Y397-phosphorylation in the residual A673 xenograft tumor cells after treatment with Y15. **C,** Schematic diagram of how EWSR1-FLI1-dependent expression of Ezrin may contribute to SRC-independent autophosphorylation of FAK on tyrosine 397 as previously described. The phosphorylation of FAK impairs apoptosis and anoikis and enhances focal adhesion formation and migratory capacity of EwS cell, but can effectively be targeted by application of Y15.

In summary, the findings presented here lead to a model where *EWSR1-ETS*-dependent expression of Ezrin leads to SRC-independent autophosphorylation of FAK on tyrosine 397 that impairs apoptosis/anoikis and enhances focal adhesion formation as well as Rho-dependent cell migration of EwS cells (Fig. 3 C). This pathway can be effectively targeted by application of the FAK inhibitor Y15.

Our results show that key tumorigenic properties of EwS cells depend on Y397-autophosphorylation of FAK, further underlining the crucial role of FA homeostasis in cancer cells, ^30^ and a possible role of FAK inhibitors in treatment of EwS. The effects of FAK inhibition were independent from the underlying *EWSR1-ETS* fusion gene. It is well conceivable that autophosphorylation of FAK depends on the structural integrity of the FA protein complex, whose components, such as Ezrin, underlie direct transcriptional control by the E/F oncogene. While further studies should aim at deciphering the exact way how expression of the oncogenic transcription factor affects FA protein composition in EwS cells, the present work clearly identified FAK autophosphorylation as a potential therapeutic target affecting metastatic dissemination in EwS.

## MATERIALS AND METHODS

### DOX-inducible knockdown and *in vivo* xenografts

A673/TR/shEF1 cells^2^, which contain a doxycycline (DOX)-inducible shRNA against *EWSR1-FLI1*, were injected subcutaneously in the flanks of immunocompromised NSG (NOD/scid/gamma) mice. When tumors reached an average volume of 180 mm^3^, mice were randomized and either received 2 mg/ml DOX (Sigma) and 5% sucrose in the drinking water (DOX +) or only 5% sucrose (DOX −). Mice were sacrificed after 96h, and tumors were isolated for RNA and histological analysis. RNA was extracted using the ReliaPrep miRNA Cell and Tissue Miniprep System (Promega). Knockdown of *EWSR1-FLI1* was confirmed by qRT-PCR^11^. The transcriptome of each tumor (*n*=3 for DOX+ and DOX−) was profiled on Affymetrix Clariom D arrays (RIN>9). Microarray data were normalized on gene level using Signal Space Transformation Robust Multi-Chip Average (SST-RMA) and Affymetrix CDF. Three FFPE cores (1 mm) were taken from each xenograft tumor to create a tissue microarray (TMA). Animal experiments were conducted in accordance with the recommendations of the European Community (86/609/EEC), the Government of Upper Bavaria (Germany), and UKCCCR (guidelines for the welfare and use of animals in cancer research).

### Immunohistochemistry of EwS patient samples

Immunohistochemistry was performed on TMA slides containing at least two representative cores (1 mm) derived from a FFPE tissue samples from 97 EwS patients from the cooperative Ewing sarcoma study (CESS) group with full clinical and follow-up information; clinico-pathological data are given in Supplementary Table 1. TMAs were stained using anti-FAK/phospho-FAK/Ezrin antibodies (1:1000; Cell signaling, Frankfurt, Germany). Immunohistochemical staining was graded as strong (2; intense staining in ≥50% of tumor cells), moderate (1; intermediate to intense staining in 1-49% of tumor cells), and negative (0, no staining or staining in <1% of tumor cells). Software-based quantification was performed with the reciprocal intensity method (ImageJ, NIH, Maryland, USA)^21^.

### Cell lines and cell-based analyses

A673, TC-32 (EWSR1-FLI1 translocated) and CADO-ES1 (EWSR1-ERG translocated) cells have been described before^14, 18, 19, 24^. Cells were grown in DMEM with 10% fetal bovine serum (FBS) and RPMI medium with 15% FBS, respectively. All cell lines were authenticated using STR analysis (data not shown). For cultivation under adhesion-free conditions, we used Softwell AF adhesion-free hydrogel plates (#SW6-AF; Matrigen Technologies, USA).

#### Western immunoblotting

Western blotting was performed using a routine protocol after cell lysis in RIPA buffer including protease and phosphatase inhibitors (#9806 and #5872 Cell signaling, Frankfurt, Germany). Grayscale intensity values were normalized to internal positive controls and measured using ImageJ software (NIH, Maryland, USA).

#### siRNA knockdown

A673, TC-32 and CADO-ES1 Ewing sarcoma cells were grown in 25 cm^2^ cell culture flasks (medium supplemented with 2% FBS) and transfected with indicated siRNA (25 pmol; cell density of 50%) using Lipofectamine RNAiMAX (Life Technologies). A set of pre-validated short interfering RNA (Silencer Select siRNA, Life Technologies) targeting EZR (Entrez Gene ID 7430) was used: #2 (ID S14796) = sense: 5□-GGCUUUCCUUGGAGUGAAAtt-3□; antisense: 5□-UUUCACUCCAAGGAAAGCCaa-3□ and #3 (ID S14797) = sense: 5□- GGAAUCAACUAUUUCGAGAtt -3□; antisense: 5□- UUUCACUCCAAGGAAAGCCaa-3□. Non-targeting control siRNA (BLOCK-iT Alexa Fluor Red Fluorescent Control, Life Technologies) was included to screen for unspecific off-target effects. After incubation for 48-72 hours, siRNA-transfected cells were lysed and knockdown efficiency was documented by Western immunoblotting.

#### xCelligence system

The xCelligence system (OLS, Bremen, Germany) was used for real-time, label-free monitoring of cell health and behavior. Cells were seeded on E-Plate 16 (cell proliferation and adhesion) or into the upper chamber of CIM-Plate 16 (cell migration). Cell proliferation, morphology change, and attachment quality was measured using electrical impedance; cell migration was monitored using the electronically integrated Boyden chamber of CIM-Plate 16 with 10% FBS as chemoattractant in the lower chamber. All experiments were performed in quadruplicate using both 20,000 and 40,000 cells for each experimental condition.

#### Immunofluorescence (IF) microscopy and focal adhesion (FA) quantification and measurement

IF staining was performed as previously described^17^. The number of FAs was quantified using ImageJ (NIH, Maryland, USA) following a published protocol with slight modifications^23^. In short, (1) 30 randomly selected cells were photographed, (2) channels were split, (2) outlines of Paxillin-positive FAs were detected, (3) objects meeting the predefined criteria for FAs (size, circularity) were counted automatically for each cell. The number of FAs was normalized to the respective cell area. FA diameter was measured using the measurement tool of ImageJ.

#### ApoTox-Glo^®^ assay

The ApoTox-Glo^®^ Assay (Promega, Mannheim, Germany) was used following the manufacturer’s protocol. A673 EwS cells were treated with Y15 (10 μmol/l) or volume-adapted concentrations of DMSO for 24h. Free AFC as a marker of viable cells was determined by measurement of fluorescence at 400 nm excitation/505 nm emission wavelengths; free R110 as a marker of necrotic cells was measured at 485 nm excitation/520 nm emission. Thirty minutes after adding Caspase-Glo^®^ 3/7-reagent, the release of aminoluciferin was measured using a luminometer. All tests were performed in triplicate and normalized to control (background), vehicle (DMSO) and Y15 compound controls.

#### Rho activity assay

Rho activity was assessed using the Rho Activity assay from Cell Signaling (#8820, Cell Signaling, Frankfurt, Germany). GTPase activity was measured based on the ability of the GTP-bound (active) form of Rho to bind Rhotekin-RBD fusion protein; this was then immunoprecipitated with glutathione resin. The level of Rho as detected in subsequent Western immunoblotting correlates with its activation state.

### Reagents

Y15 (1,2,4,5-Benzenetetraamine tetrahydrochloride; FAK Inhibitor 14, Cat. No. 3414) was obtained in pharmaceutical purity from Tocris Bioscience (Bristol, UK).

### Chorioallantoic membrane (CAM) model

CAM assays were performed as previously described^13^. Fertilized eggs of White Leghorn chickens were incubated at 37°C with 60% relative humidity and prepared for implantation on day 4 of incubation. Each egg was washed with warm 70% ethanol and a hole was drilled through the pointed pole of the shell. The following day, the chorioallantoic membrane was exposed by peeling a 1.5–2.0 cm window in the shell. This window was covered with tape and the incubation continued. On day 8 of incubation, DMSO/Y15-treated A673 EwS cells were dissolved in Matrigel and introduced in the CAM. Tumor growth was assessed macroscopically during the following days. After 5 days, tumors were harvested, measured and weighed, fixed in 5% PFA and processed for histopathological examination with H/E, FAK/pFAK (Y397) and Ezrin staining as described above.

### Statistical analyses

Survival analyses (Kaplan-Meier method) for the patient-derived sample cohorts were done with SPSS (IBM, Mannheim, Germany). All statistical analyses for the *in vitro* data were performed using GraphPad software (GraphPad, LaJolla, USA). We used Chi-Square/Fisher’s exact test for the comparison of categorical variables, while student’s t-test/ANOVA followed by Tukey’s multiple comparison test was performed to compare continuous variables between two groups/more than two groups, respectively. A *p* value <0.05 was regarded as statistically significant.

---

**Supplementary Figure 1. A,** Detection of EWSR1-FLI1 and EWSR1-ERG fusion proteins in TC-32, A673 and CADO-ES1 cells. **B,** siRNA-based knockdown of Ezrin in TC-32 (48h) and CADO-ES1 (48h and 72h, respectively). **C,** Compared to TC-32 and CADO-ES1, A673 showed the highest proliferative activity and migratory potential in real-time proliferation and migration assays using the xCelligence system. **D and E,** Software-based image analysis (ImageJ) showed numerous Paxillin-positive focal adhesions (FA) in the investigated EwS cell lines, with the highest number of FAs observed in A673 and CADO-ES1. **F,** FA protein expression in EwS cells. Strong expression and Y397-phosphorylation of FAK in all investigated EwS cell lines, while FAK Y576/577 expression was only barely detectable. Y118-phosphorylation of Paxillin was present only in CADO-ES1 cells

