## Supplemental Table 1 for "Focal adhesion kinase confers pro-migratory and anti-apoptotic properties and is a potential therapeutic target in Ewing sarcoma"

**Supplementary Table 1**

**Clinico-pathological data**

|  | **total** | **%** |
| --- | --- | --- |
| **No. of patients** | 97 | 100 |
| **Sex**  Male  Female | 62  35 | 64  36 |
| **Age at diagnosis (years)**  Mean  Median  Range | 19.8  17.0  0-68 |  |
| **Tumor volume**  <200 ml  ≥200 ml  n.a. | 48  44  5 | 49  45  6 |
| **Primary metastases**  present absent | 37  60 | 38  62 |
| **FAK staining intensity**  Strong  Moderate  Negative  n.a. | 92  1  0  4 | 95  1  0  4 |
| **pFAK (Y397) staining intensity**  Strong  Moderate  Negative  n.a. | 41  35  14  7 | 42  36  14  7 |
| **Ezrin staining intensity**  Strong  Moderate  Negative  n.a. | 43  34  13  7 | 44  35  13  7 |
| **Response to CTX**  Good  Poor  n.a. | 47  12  38 | 48  12  39 |
| **Time to first relapse (months)**  Mean  Median  Range  Complete response | 29.5  17.3  4.3-170.3  42 | 43 |
| **Type of first relapse**  Local  Systemic  Combined  Complete response | 4  41  10  42 | 4  42  10  43 |
| **OS (months)**  Mean  Median  Range | 68.8  53.5  8.6-222.1 |  |
| **Deaths** | 50 | 52 |
