## Supplementary figures and images for "Focal adhesion kinase confers pro-migratory and anti-apoptotic properties and is a potential therapeutic target in Ewing sarcoma"

### Supplemental Data 1

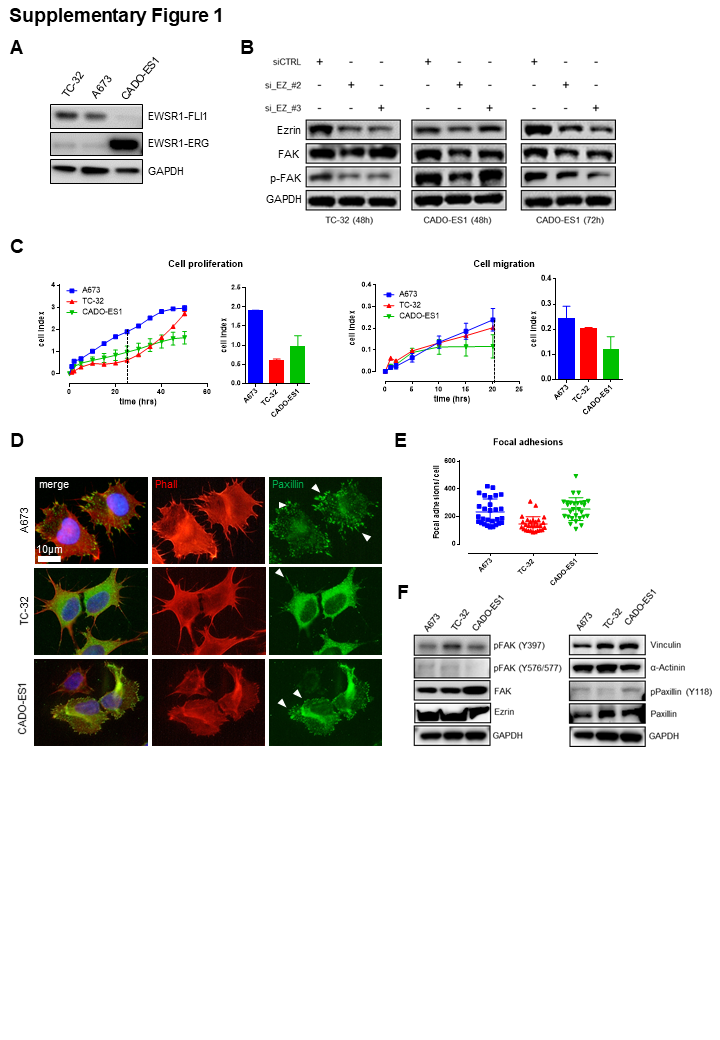
